# Targeting Macrophage Folate Receptor-β for Diagnosis and Treatment of Autoimmune Myocarditis

**DOI:** 10.1101/2025.11.02.686077

**Authors:** Arghavan Jahandideh, Erika Atencio Herre, Wail Nammas, Jenni Virta, Heidi Liljenbäck, Jesse Ponkamo, Putri Andriana, Xiaoqing Zhuang, Johan Rajander, Madduri Srinivasarao, Philip S. Low, Yingjuan June Lu, Xiang-Guo Li, Juhani Knuuti, Anne Roivainen, Antti Saraste

## Abstract

We investigated folate receptor-β (FR-β) expressed on activated macrophages as a therapeutic and imaging target in autoimmune myocarditis (AIM). Treatment with a folate-aminopterin conjugate (EC2319) targeting FR-β-positive macrophages significantly reduced macrophage infiltration compared to saline (5.23 ± 2.84% vs. 1.24 ± 0.66%, *P* < 0.001) in rat experimental AIM. After immunization, FR-β targeting positron emission tomography (PET) tracer [^18^F]FOL detected myocardial inflammatory lesions and revealed reduction in inflammation following EC2319 treatment relative to saline or tumor necrosis factor-alpha inhibitor Etanercept [standardized uptake value (SUV) 1.07 ± 0.22 vs. 2.38 ± 0.90 vs. 2.59 ± 0.56, *P* < 0.001]. [^18^F]FOL PET signal correlated with myocardial CD68-positive macrophage content (*r* = 0.720, *P* < 0.001). These findings demonstrate that EC2319 attenuates myocardial inflammation in AIM and underscore the pathogenic role of FR-β-positive macrophages. [^18^F]FOL PET provides a noninvasive means of detecting myocardial inflammatory activity and therapeutic response.

**Graphical abstract:** 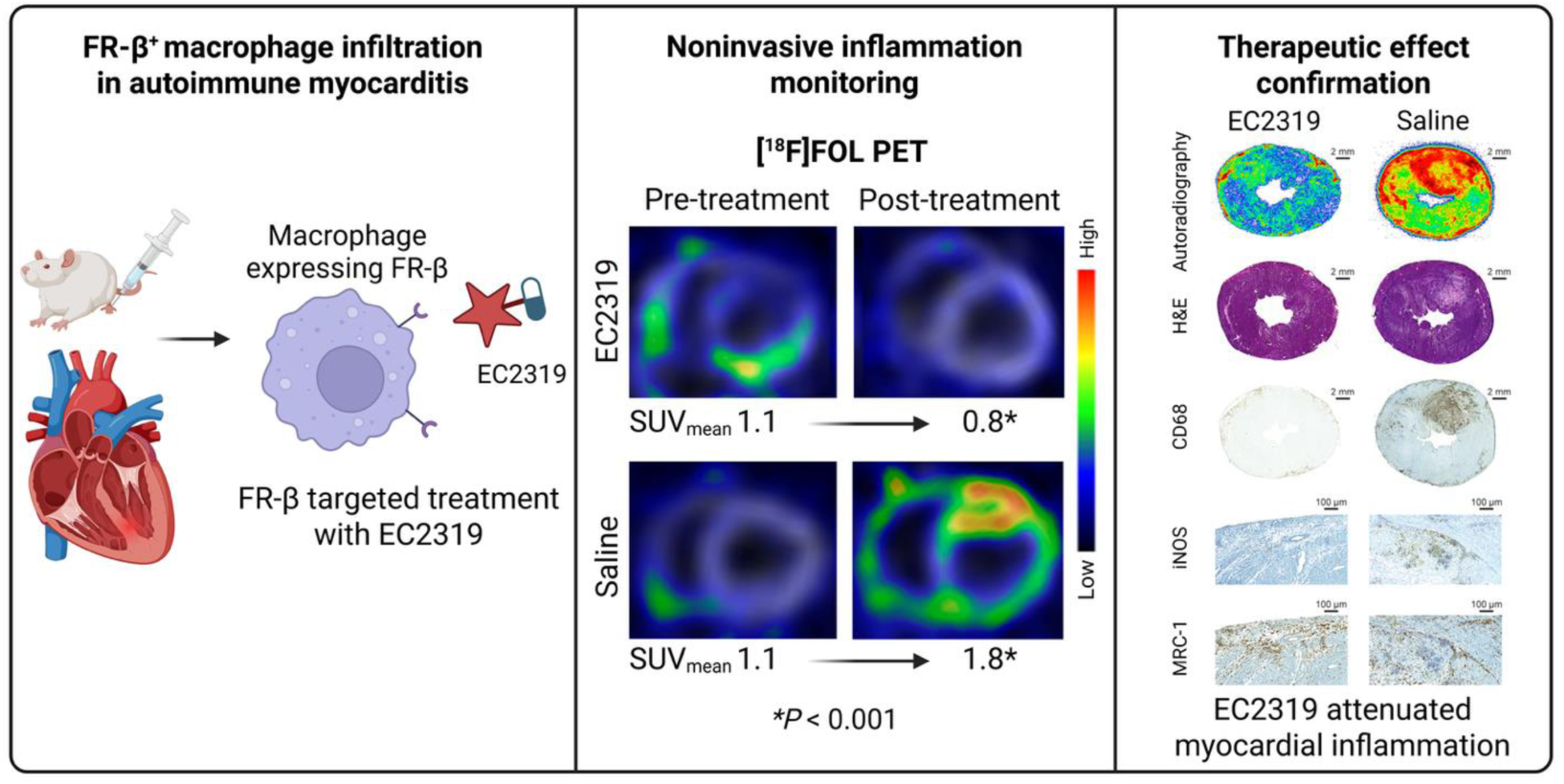

## Introduction

Myocarditis is an inflammatory disease of the heart characterized by myocardial infiltration of inflammatory cells and myocyte necrosis (1). It may result from infections, cardiotoxic agents, or autoimmune conditions, such as cardiac sarcoidosis (1–3). Clinical manifestations include conduction abnormalities, ventricular arrhythmias, and inflammatory cardiomyopathy, defined as myocarditis associated with impaired cardiac function (1–3). Advanced non-invasive imaging modalities, including cardiac magnetic resonance imaging and positron emission tomography (PET) using 2-deoxy-2-[^18^F]fluoro-*D*-glucose ([^18^F]FDG), are instrumental in characterizing myocardial abnormalities in myocarditis (2, 3).

Activated macrophages express the functional folate receptor-β (FR-β), a glycosylphosphatidylinositol-anchored membrane protein that binds folic acid and folate-conjugated small molecules with high affinity, internalizing them via endocytosis (4–7). FR-β expression is upregulated in inflammatory conditions, making it a validated target for imaging and therapeutic interventions in inflammatory diseases (5–10). In early clinical studies, folate receptor-specific radiotracers have successfully identified activated macrophages in inflamed joints (11–13). We previously demonstrated that FR-β is expressed in cardiac inflammatory lesions in both an experimental autoimmune myocarditis (AIM) rat model and in patients with cardiac sarcoidosis (10). In experimental AIM, PET imaging with aluminum fluoride-18-labeled 1,4,7-triazacyclononane-*N*,*N*′,*N*″-triacetic acid-conjugated folate ([^18^F]FOL) effectively detected FR-β-positive macrophage infiltration in the myocardium, achieving high signal-to-noise ratio (10). However, whether FR-β can be exploited as a therapeutic target and biomarker for monitoring response to anti-inflammatory treatment in myocarditis remains unknown.

Immunosuppressive therapy aims to suppress active inflammation and prevent development of irreversible myocardial injury and heart failure in AIM (2, 3). Aminopterin is a dihydrofolate reductase inhibitor with potent anti-inflammatory activity similar to methotrexate (14, 15), which is used as a second-line immunosuppressant in treatment of cardiac sarcoidosis (2, 3). A novel folate-aminopterin conjugate, EC2319, has been shown to resolve inflammation in various disease models by selectively targeting FR-β-expressing monocytes and macrophages (9, 14, 15).

The aim of this study was to evaluate the therapeutic potential of FR-β-targeted antifolate therapy using EC2319 in a rat model of AIM, and to assess [^18^F]FOL PET as a non-invasive tool for monitoring treatment response. We first investigated whether EC2319 administration during immunization could prevent the development of myocarditis and cardiac dysfunction. In a subsequent intervention sub-study, we compared the therapeutic effects of EC2319, the tumor necrosis factor-alpha (TNF-α) inhibitor Etanercept, and saline in rats with established myocarditis verified by [^18^F]FOL PET. Treatment response was evaluated by PET imaging and histological assessment of myocardial inflammation.

## Results

### Effects of EC2319 on the severity of autoimmune myocarditis

In the prevention sub-study, all 24 rats successfully completed AIM induction, treatment, and follow-up until day 21 after the first immunization. Histological examination revealed focal myocardial inflammatory lesions characterized by dense infiltration of CD68-positive macrophages and associated myocyte loss. In the saline-treated group (*n* = 12), inflammatory lesions were observed in 92% of rats (Figure 1), predominantly in the left ventricle (LV), and occasionally in the right ventricle (RV) myocardium. Echocardiography showed pericardial effusion of varying severity in 5 of 12 rats and a significant increase in LV end-systolic volume (LVESV), along with a corresponding decrease in LV ejection fraction (LVEF) from baseline to the end of study (Table 1).

**Figure 1.**
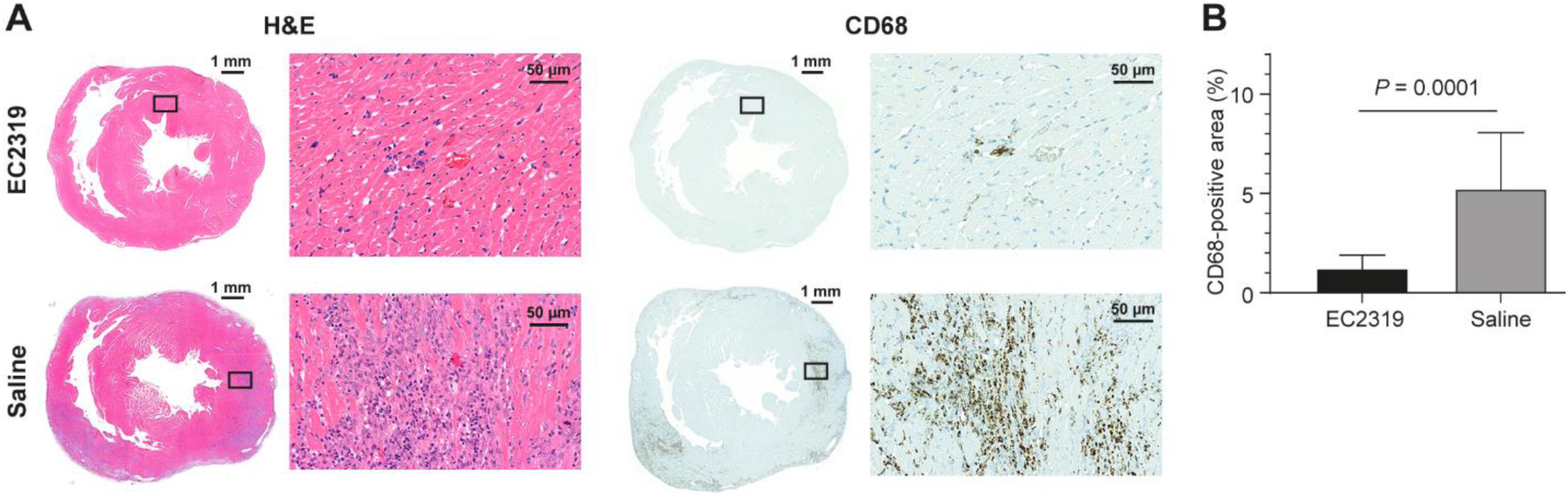
Effect of EC2319 on myocardial inflammation in the prevention sub-study. (**A**) Representative images of hematoxylin-eosin (H&E) staining and immunohistochemistry for CD68-positive macrophages in rat hearts collected 21 days after the first immunization. Rats were administered EC2319 or saline throughout the induction of myocarditis. (**B**) Quantification of CD68-positive macrophages in myocardial tissue on Day 21 of the prevention sub-study.

**Table 1.** Echocardiographic parameters before and after treatment with EC2319, TNF-α inhibitor (Etanercept), or saline

|  | EC2319 | Saline | Etanercept | *P*<br>value* | *P*<br>value** | *P*<br>value*** | *P*<br>value**** |
| --- | --- | --- | --- | --- | --- | --- | --- |
| **EC2319 treatment during immunization** |  |  |  |  |  |  |  |
| **Before immunization** |  |  |  |  |  |  |  |
| Heart rate | 386.8 ± 86.7 | 384.1 ± 23.5 | NA | 0.551 | 1.000 | 0.025 | NA |
| LVEF (%) | 62.6 ± 10.2 | 64.6 ± 11.5 | NA | 0.651 | 0.345 | 0.027 | NA |
| LVEDV (μL) | 520.5 ± 150.4 | 527.7 ± 70.7 | NA | 0.883 | 0.520 | 0.739 | NA |
| LVESV (μL) | 193.8 ± 87.4 | 188.1 ± 71.0 | NA | 0.862 | 0.511 | 0.047 | NA |
| **After treatment** |  |  |  |  |  |  |  |
| Heart rate | 386.8 ± 38.0 | 356.0 ± 35.5 | NA | 0.114 | | | |
| LVEF (%) | 57.1 ± 15.4 | 50.0 ± 12.3 | NA | 0.225 | | | |
| LVEDV (μL) | 559.8 ± 130.5 | 519.0 ± 106.1 | NA | 0.410 | | | |
| LVESV (μL) | 221.3 ± 96.5 | 257.0 ± 77.1 | NA | 0.327 | | | |
| **EC2319 or Etanercept treatment after immunization** |  |  |  |  |  |  |  |
| **After immunization** |  |  |  |  |  |  |  |
| Heart rate | 395.5 ± 25.0 | 364.2 ± 19.5 | 385.0 ± 24.0 | 0.052 | 0.330 | 0.034 | 0.854 |
| LVEF (%) | 70.0 ± 9.8 | 67.1 ± 11.0 | 71.7 ± 7.0 | 0.669 | 0.616 | 0.243 | 0.811 |
| LVEDV (μL) | 405.3 ± 84.6 | 405.1 ± 128.5 | 414.2 ± 128.6 | 0.988 | 0.003 | 0.049 | 0.859 |
| LVESV (μL) | 119.0 ± 38.0 | 145.4 ± 76.8 | 123.9 ± 62.1 | 0.810 | 0.059 | 0.076 | 0.839 |
| **After treatment** |  |  |  |  |  |  |  |
| Heart rate | 379.6 ± 23.5 | 388.7 ± 20.0 | 381.5 ± 38.8 | 0.606 | | | |
| LVEF (%) | 68.2 ± 6.7 | 61.9 ± 9.9 | 70.1 ± 14.6 | 0.240 | | | |
| LVEDV (μL) | 477.3 ± 72.0 | 482.7 ± 88.5 | 427.3 ± 82.1 | 0.358 | | | |
| LVESV (μL) | 149.7 ± 27.5 | 182.6 ± 56.0 | 118.5 ± 35.2 | 0.017 | | | |

Results expressed as mean ± SD. Non-paired *t*-tests or ANOVA were done comparing effects of treatments; paired *t*-tests were done to compare results at baseline and end of study. LVEDV=left ventricular end diastolic volume, LVEF=left ventricular ejection fraction, LVESV=left ventricular end systolic volume, NA=not available, \*=between treatments, \*\*=EC2319 before vs. after treatment, \*\*\*=saline before vs. after treatment, \*\*\*\*=Etanercept before vs. after treatment

Treatment with EC2319 (*n* = 12), administered during immunization, was well tolerated. Rats receiving EC2319 demonstrated greater weight gain from day 0 to study completion compared to saline-treated controls (38.9 ± 17.8 g vs. 17.4 ± 20.5 g; *P* = 0.010). Histologically, myocardial inflammatory lesions were observed in only 25% of EC2319-treated rats. Quantitative analysis showed a significantly lower percentage area of CD68-positive macrophage infiltration in the myocardium of EC2319 group compared to the saline group (1.2 ± 0.7 % vs. 5.2 ± 2.8 %; *P* < 0.001). Pericardial effusion was detected in 3 of 12 rats treated with EC2319. Notably, heart rate, LV volumes, and LVEF remained unchanged from baseline in the EC2319 group. By the end of the study, there were no significant differences in LV volumes or LVEF between EC2319-and saline-treated animals (Table 1).

### Effects of EC2319 on myocardial inflammation assessed by [^18^F]FOL PET

In the intervention sub-study, 22 of the 37 scanned rats demonstrated visually detectable myocardial uptake of [^18^F]FOL on PET imaging consistent with myocardial inflammation after completion of immunization on day 13. These rats were randomized to receive EC2319 (*n* = 8), saline (*n* = 8), or Etanercept (*n* = 6). All animals completed treatment and follow-up through day 26. Weight gain from day 13 to the end of the study was comparable across groups (EC2319: 33.6 ± 12.8 g; saline: 31.6 ± 28.0 g; Etanercept: 38.0 ± 29.0 g; *P* = 0.884).

[^18^F]FOL PET imaging revealed focal myocardial uptake before treatment (day 13) and after treatment (day 26), with uptake co-localizing with histological inflammatory lesions and uptake of [^18^F]FOL seen by autoradiography in tissue sections (Figures 2 and 3). In PET images, [^18^F]FOL uptake was quantified as standardized uptake value (SUV) of segment with the highest uptake (SUV_max_) or mean SUV across the LV myocardium (SUV_mean_) using 17-segment polar maps of the LV myocardium. Representative [^18^F]FOL PET images and polar maps as well as SUVs before and after treatments are shown in Figure 2.

**Figure 2.**
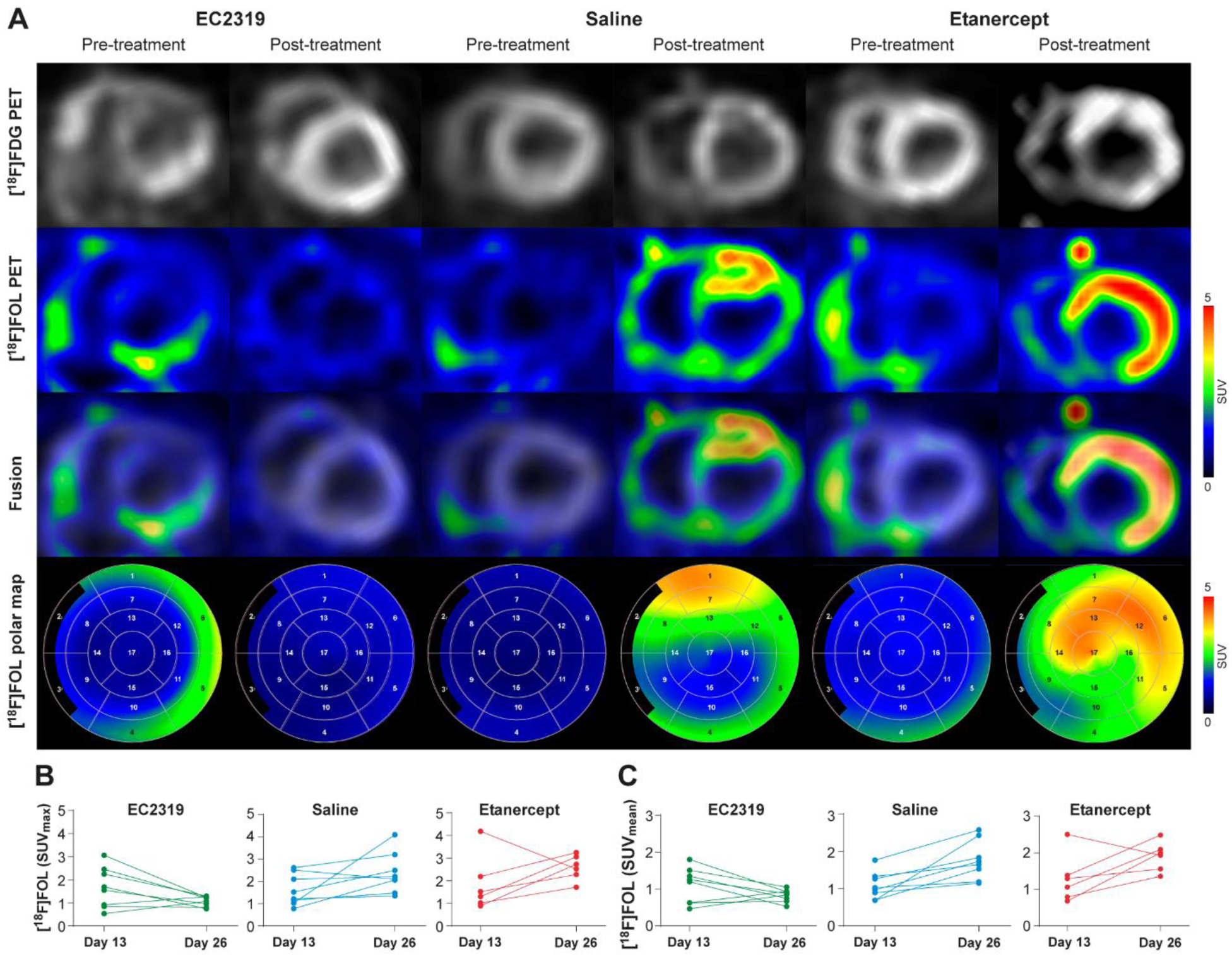
Assessment of myocardial inflammation by [¹⁸F]FOL PET imaging before and after treatment. (**A**) Representative short axis [^18^F]FOL PET images and corresponding polar maps of the left ventricle (LV) of rat hearts at baseline (Day 13, pre-treatment) and post-treatment (Day 26). Images show the presence of uptake of [^18^F]FOL in all rats pre-treatment, and more extensive uptake in rats administered with saline or Etanercept than EC2319 post-treatment. Note that histology and autoradiographs of [^18^F]FOL uptake in tissue sections corresponding to PET images shown in Figure 3 demonstrate co-localization of [^18^F]FOL uptake with inflammatory lesions. (**B)** Changes in [^18^F]FOL uptake (standardized uptake value, SUV) in the most inflamed myocardial segment post-treatment (SUV_max_) and (**C**) mean [¹⁸F]FOL uptake across the entire LV myocardium (SUV_mean_).

**Figure 3.**
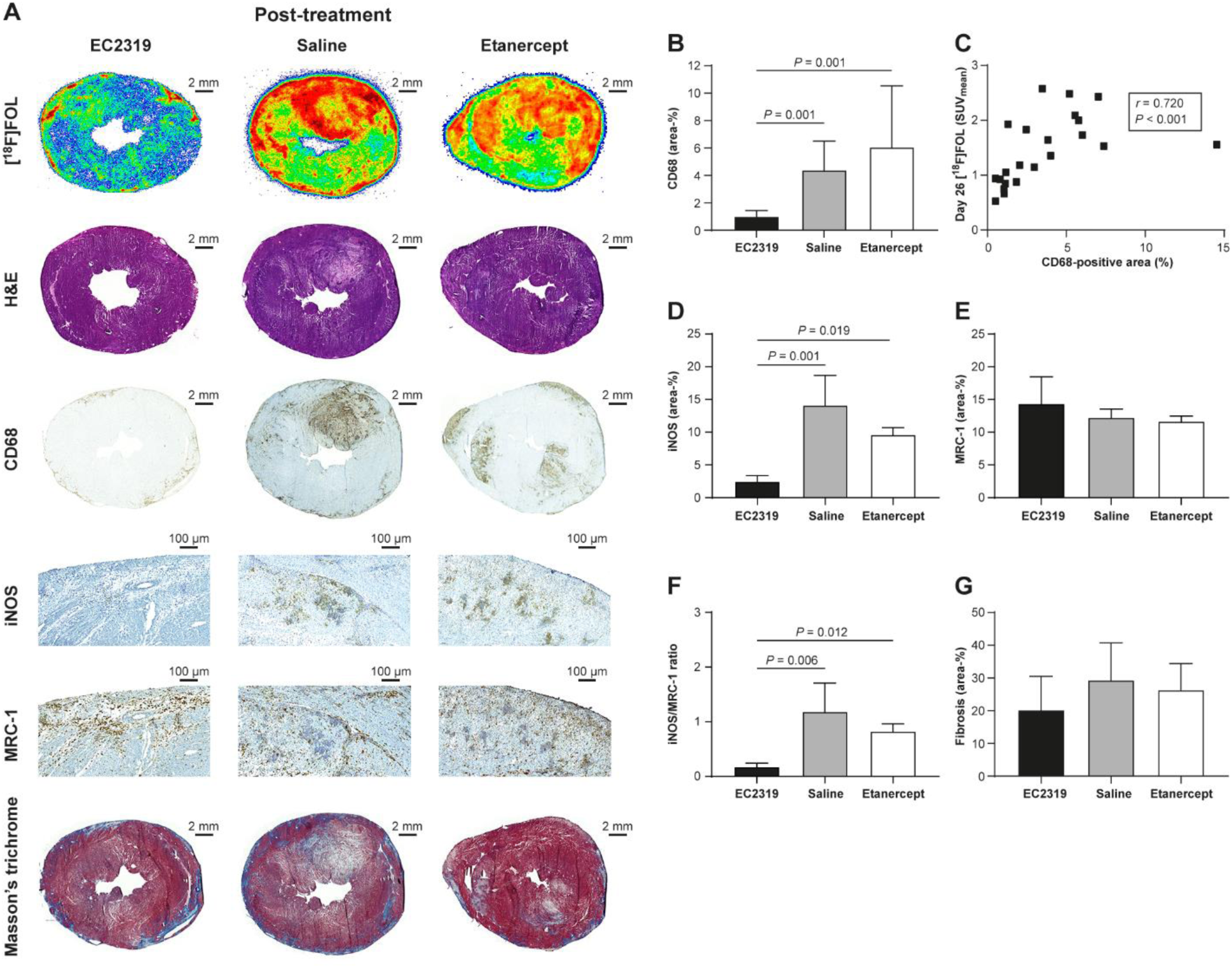
Histological and molecular characterization of myocardial inflammation and fibrosis following treatment. (**A**) Representative [^18^F]FOL autoradiographs and corresponding hematoxylin and eosin (H&E) and immunohistochemical stainings of myocarditis rat hearts post-treatment. (**B**) Quantification of CD68-positive macrophages and (**C**) correlation with myocardial [^18^F]FOL uptake on Day 26. Spearman correlation coefficient was used to analyze linear correlations between CD68-positive area and myocardial [^18^F]FOL uptake on Day 26. (**D**) Quantifications of M1-polarized macrophages (iNOS-positive), (**E**) M2-polarized macrophages (MRC-1-positive), and (**F**) M1/M2 macrophage ratios in inflammatory lesions based on immunostaining. (**G**) Quantification of fibrotic area using Masson’s trichrome staining. *n* = 6-8 per group for B-C; *n* = 3 per group for D-F; *n* = 5-7 per group for G. Statistical comparisons of normally distributed data were performed using one-way ANOVA followed by post hoc tests: Tukey’s test (for homogeneous variances) and Games–Howell test (for non-homogeneous variances) to compare different treatments. Non-normally distributed continuous data are presented as median (interquartile range, IQR) and compared using the Kruskal–Wallis test, followed by pairwise comparisons with Bonferroni-adjusted *p*-values.

At baseline, myocardial [^18^F]FOL uptake measured as either SUV of segment with the highest uptake (SUV_max_) or mean SUV in the LV (SUV_mean_) was similar across all treatment groups (SUV_max_; *P* = 0.716; SUV_mean_: *P* = 0.893; Table 2, Figure 2). After treatment, EC2319-treated rats demonstrated significantly lower myocardial [^18^F]FOL uptake compared to both saline-and Etanercept-treated groups (Table 2). Specifically, SUV_max_ was significantly lower in the EC2319 group compared to saline (*P* = 0.002) and Etanercept (*P* < 0.001). Similarly, SUV_mean_ in the LV was significantly lower after EC2319 treatment compared to both saline and Etanercept groups (*P* < 0.001 for both). Compared with baseline, SUV_mean_ decreased by 53.8 ± 23.7% (*P* = 0.134) in EC2319 treated rats, whereas SUV_mean_ increased in both saline (30.6 ± 7.9%; *P* = 0.006 vs. EC2319) and Etanercept (31.7 ± 16.7%; *P* = 0.010 vs. EC2319) groups (Table 2, Figure 2). Furthermore, myocardial segment with the highest SUV before treatment showed significantly lower [^18^F]FOL uptake after EC2319 treatment [median 0.8 (interquartile range, IQR 0.4)] compared to both saline [median 1.4 (IQR 0.9); *P* = 0.002] and Etanercept [median 2.1 (IQR 1.6); *P* < 0.001] treatments.

**Table 2.** [^18^F]FOL PET imaging of myocardial uptake in immunized rats before and after treatments

|  | EC2319 | Saline | Etanercept | *P* value* | *P* value** | *P* value*** | *P* value**** |
| --- | --- | --- | --- | --- | --- | --- | --- |
| **Before treatment** |  |  |  |  |  |  |  |
| SUV<sub>mean</sub> | 1.1 ± 0.5 | 1.1 ± 0.4 | 1.3 ± 0.5 | 0.716 | 0.994 | 0.781 | 0.722 |
| SUV<sub>max</sub> | 1.7 ± 0.9 | 1.7 ± 0.7 | 1.9 ± 1.2 | 0.893 | 0.996 | 0.926 | 0.892 |
| **After treatment** |  |  |  |  |  |  |  |
| SUV<sub>mean</sub> | 0.8 ± 0.2 | 1.8 ± 0.5 | 1.9 ± 0.4 | < 0.001 | < 0.001 | < 0.001 | 0.777 |
| SUV<sub>max</sub> | 1.1 ± 0.2 | 2.4 ± 0.9 | 2.6 ± 0.6 | < 0.001 | 0.002 | < 0.001 | 0.812 |

Results expressed as SUVs (mean ± SD). One-way ANOVA and post hoc tests were done comparing effects of different treatments. \* = Overall, \*\* = EC2319 vs. Saline, \*\*\* = EC2319 vs. Etanercept, \*\*\*\* = Saline vs. Etanercept.

### Ex vivo validation and histological assessment of inflammatory activity

Consistent with in vivo PET findings, ex vivo gamma counting demonstrated significantly higher [^18^F]FOL radioactivity in the hearts of rats treated with saline or Etanercept compared to those treated with EC2319 (SUV saline: 2.7 ± 0.4, Etanercept: 3.6 ± 0.6, and EC2319: 1.1 ± 0.1; *P* < 0.001 for both). Biodistribution analysis (Supplemental Table 1) further revealed lower [^18^F]FOL radioactivity in the spleens of EC2319-treated rats compared to saline- or Etanercept-treated (SUV 2.6 ± 0.3 vs. 5.9 ± 1.1 vs. 9.8 ± 0.6; *P* < 0.001). A similar pattern was observed in the liver, thymus and skull bone (Extended Data Table 1).

Histological analysis revealed markedly reduced myocardial inflammation in EC2319-treated rats compared to both control groups (*P* < 0.001). The stained area fraction for CD68-positive macrophage immunohistochemistry was significantly lower in the EC2319 group than in saline-treated [median 1.1 (IQR 0.6) % vs. 3.7 (IQR 4.2) %; *P* = 0.001) or Etanercept-treated rats (median 5.4 (IQR 4.6) %; *P* = 0.001; Figure 3, A and B). There was a significant correlation between [^18^F]FOL uptake (SUV_mean_) and stained area fraction for CD68-positive macrophages (*r* = 0.720; *P* < 0.001, Figure 3C).

In inflammatory lesions, the stained area fraction for inducible nitric oxide synthase (iNOS), a marker of pro-inflammatory M1 macrophages, was significantly lower in the EC2319 group (2.6 ± 0.8 %) compared to saline (14.1 ± 3.7 %; *P* = 0.001) and Etanercept groups (9.6 ± 0.9 %; *P* = 0.019; Figure 3, A and D). In contrast, stained area fraction for mannose receptor C type 1 (MRC-1, also known as CD206), a marker of anti-inflammatory M2 macrophages, did not differ between inflammatory lesions of EC2319-treated rats and those treated with saline (14.5 ± 4.1 % vs. 12.3 ± 1.0 %; *P* = 0.602) or Etanercept (11.7 ± 0.6 %; *P* = 0.456; Figure 3, A and E). Consequently, the iNOS/MRC-1 ratio was lower in inflammatory lesions of EC2319-treated rats compared to those treated with saline (0.2 ± 0.1 % vs. 1.2 ± 0.4 %; *P* = 0.006) or Etanercept (0.8 ± 0.1 %; *P* = 0.012; Figure 3F), indicating a relative shift toward reduced pro-inflammatory macrophage predominance. Consistent with this, Figure 4 shows co-localization of CD68-immunofluorescence predominantly with iNOS, which was lower after treatment with EC2319 than saline or Etanercept.

**Figure 4.**
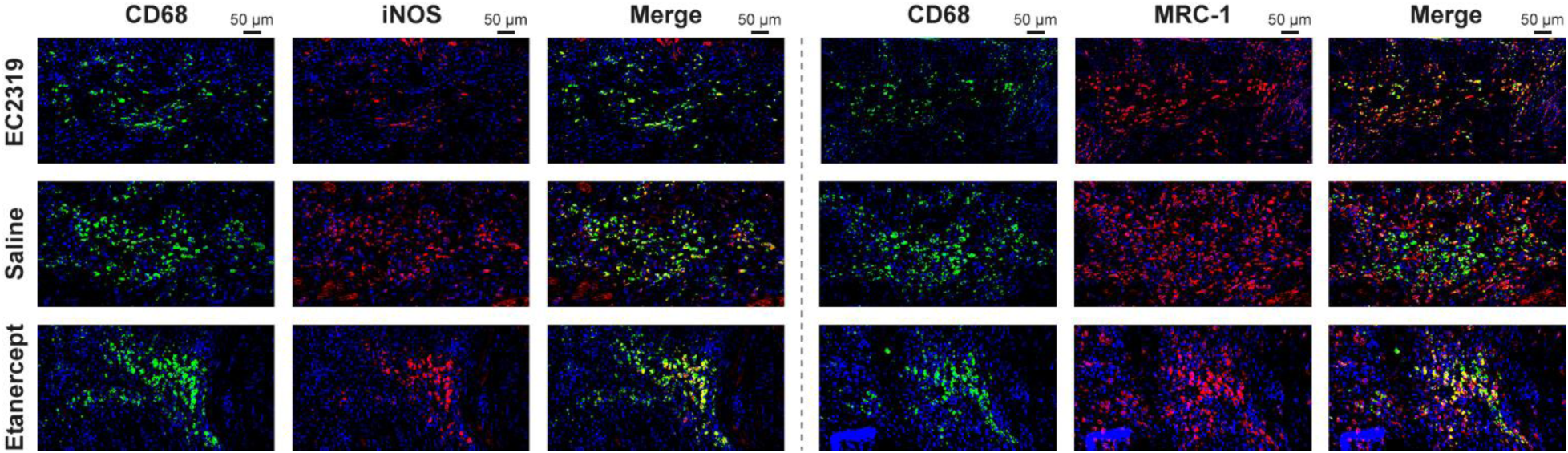
Immunofluorescence staining of macrophage polarization markers after treatment. There was strong co-localization of CD68 and iNOS, a marker of pro-inflammatory M1 macrophages, whereas there was only partial co-localization of CD68 and MRC-1, a marker of anti-inflammatory M2 macrophages. Number of iNOS-positive macrophages is markedly lower after treatment with EC2319 than saline or Etanercept, consistent with a shift toward an anti-inflammatory macrophage profile.

Masson’s trichrome staining revealed comparable levels of myocardial fibrosis across treatment groups. The fibrotic area was not significantly different between EC2319-treated rats (20.3 ± 10.2 %) and those treated with saline (29.4 ± 11.3 %; *P* = 0.297) or Etanercept (26.4 ± 8.0 %; *P* = 0.586; Figure 3G).

After treatment, echocardiography showed no significant differences in heart rate, LVEF, or LVEDV among rats treated with EC2319, saline, or Etanercept (Table 1). LVESV was also comparable between EC2319- and saline-treated groups (*P* = 0.172).

## Discussion

In this study, we demonstrate that the FR-β-targeted folate-aminopterin conjugate EC2319 effectively attenuates myocardial inflammation in a rat model of AIM. Administration of EC2319 either during the immunization phase or after the onset of myocarditis significantly reduced macrophage-rich inflammatory lesions, whereas treatment with the TNF-α inhibitor Etanercept had no effect. Our results indicate a key role for FR-β-expressing pro-inflammatory macrophages in mediating myocardial injury in AIM. Furthermore, PET imaging with the FR-β-targeted radiotracer [^18^F]FOL enabled non-invasive monitoring of treatment response in the inflamed myocardium.

The rat AIM model is a well-established experimental model that recapitulates several pathological features of human giant cell myocarditis, including a fulminant inflammatory phase (days 14-28 after immunization) progressing to myocardial fibrosis and dilated cardiomyopathy (10, 20, 21). After the initial autoimmune activation involving autoreactive T cells and autoantibodies targeting heart antigens, monocyte-derived macrophages contribute significantly to disease progression and the formation of giant cells and granulomatous lesions – hallmarks of giant cell myocarditis and cardiac sarcoidosis^22^. Consistent with human AIM, macrophages dominate the cellular infiltrate in this model (10, 21, 22).

Activated macrophages upregulate FR-β and exhibit a pro-inflammatory phenotype, producing reactive oxygen species and TNF-α (5–7). We previously identified FR-β expression in inflammatory myocardial lesions in this AIM model, as well as in granulomatous cardiac lesions from patients with cardiac sarcoidosis (10). EC2319, a folate-aminopterin derivative, selectively targets FR-β-expressing cells and has been shown to inhibit pro-inflammatory activation of monocytes and macrophages, thereby attenuating inflammation in various experimental disease models (9, 14, 15). As the role of FR-β in AIM pathogenesis had not been previously defined, we investigated the therapeutic efficacy of FR-β-targeted treatment with EC2319 in this model.

In the prevention sub-study, EC2319 administered during immunization and continued through the period of active inflammation resulted in a marked reduction in disease severity, with >400% decrease in myocardial CD68-positive macrophages burden and inflammatory lesions in only 25% of treated rats. In untreated animals, the inflammatory process led to reduction in LVEF, whereas EC2319-treated animals maintained preserved systolic function, consistent with decreased myocardial damage. In the intervention sub-study, EC2319 given during the fulminant phase of myocarditis reduced the extent of myocardial CD68-positive macrophage infiltration to levels comparable to those seen in the prevention group and >200% lower than in rats treated with saline or Etanercept. These findings support earlier studies showing that macrophage depletion or chemokine receptor blockage can mitigate myocardial inflammation and improve cardiac function in AIM models (23–25). Our data further highlight a central role for FR-β-positive macrophages in disease pathogenesis and support FR-β-targeted anti-inflammatory therapy as a promising approach for suppressing active myocardial inflammation in this setting.

Methotrexate and aminopterin, both folate antagonists, act by inhibiting dihydrofolate reductase and other folate-dependent enzymes, thereby impairing nucleotide synthesis, modulating cytokine profiles, and increasing extracellular adenosine—all of which suppress immune cell proliferation and inflammation (9, 14). EC2319 and the related compound EC0746 represent FR-β-targeted aminopterins with approximately 40-fold improved tolerability compared to unmodified aminopterin (9, 14). These compounds exhibit selective activity against FR-β-positive inflammatory monocytes and macrophages, including inhibition of dihydrofolate reductase, cytostatic effects, and cytokine modulation (9, 14, 15, 26). Our dosing regimen was based on prior studies (9, 14, 15, 26), and to minimize competition with endogenous folate, rats were fed a folate-deficient diet (9). EC2319 was well-tolerated, with no observed toxicity or mortality during the study period, and treated rats showed greater weight gain compared to saline controls. These findings are consistent with the established safety profile of FR-β-targeted aminopterins. Methotrexate, often used in combination with other immunosuppressants, has shown benefit in treating AIM and is currently being evaluated in a clinical trial (2, 3). In experimental AIM, methotrexate treatment started at the late fulminant phase has been shown to reduce systemic inflammation and attenuate myocardial fibrosis and dysfunction (27, 28). Our findings extend these observations, demonstrating that FR-β-targeted antifolate therapy can effectively suppress active myocardial inflammation and may limit inflammation-driven myocardial injury.

We previously showed that [^18^F]FOL PET enables detection of active myocardial inflammation in experimental AIM (10). Unlike [^18^F]FDG PET, which may lack specificity due to its uptake by physiological myocardial glucose metabolism, [^18^F]FOL targets FR-β and offers improved specificity (2, 3, 29). In the present study, [^18^F]FOL PET imaging confirmed active myocardial inflammation at baseline and demonstrated treatment response. The finding that [^18^F]FOL PET revealed inflammation in 59% of rats after immunization is consistent with previous observations that roughly one third do not develop myocarditis (10), and highlights the potential of imaging to guide therapy. After treatment, a 4-day washout was implemented prior to follow-up PET to avoid interference from residual EC2319, based on its known pharmacokinetics in rats (15). Compared to saline or Etanercept, EC2319 treatment significantly reduced SUV_max_ reflecting the most intense myocardial inflammation and SUV_mean_ reflecting inflammatory burden across the LV. Furthermore, PET signal correlated with myocardial CD68-positive macrophage burden, supporting [^18^F]FOL PET as a quantitative method to assess the degree of inflammation in myocarditis. A decline in [^18^F]FOL uptake in initially active regions following treatment further supports the efficacy of EC2319 in reducing existing myocardial inflammation. These results validate [^18^F]FOL PET as a sensitive, non-invasive biomarker for detecting and monitoring myocardial inflammation and therapeutic response.

We also examined macrophage polarization states, given that multiple subsets exist in the heart (22, 30). Macrophages express FR-β upon activation by pro-inflammatory factors, and FR-β-positive macrophages both express pro-inflammatory markers and secrete pro-inflammatory mediators, such as reactive oxygen species and TNF-α (5–7). However, FR-β has been reported on both M1- and M2-like macrophages (31, 32). In our study, iNOS-positive (M1-like) macrophages were more abundant than MRC-1-positive (M2-like) cells in inflammatory lesions. EC2319 treatment led to marked reduction in iNOS-positive cells and decreased iNOS:MRC-1 ratio. This pattern is consistent with previous reports in models such as experimental autoimmune encephalomyelitis (15, 21), and with the concepts that EC2319 predominantly targets subpopulation of pro-inflammatory macrophages and modulates the balance between pro-inflammatory and anti-inflammatory macrophage subsets. Although detailed temporal dynamics of FR-β expression remain unknown, our study suggests that FR-β-targeted therapy can both prevent development of myocarditis and ameliorate ongoing inflammation. Additionally, EC2319 treatment reduced [^18^F]FOL uptake in hematopoietic organs such as spleen and bone marrow, suggesting a systemic immunomodulatory effect consistent with reduced hematopoietic activation during AIM (9, 14).

Etanercept was selected as a comparator based on reports of its potential benefit in AIM when used in combination with other immunosuppressants (2, 3, 33). The dose tested was previously shown to be effective in rodent models of rheumatoid arthritis (9). However, in our study, Etanercept alone had no impact on myocardial inflammation or [^18^F]FOL uptake. This aligns with recent evidence suggesting that TNF-α may paradoxically exert protective effects by promoting activation-induced cell death of autoreactive T cells in myocarditis (34). Similarly, clinical outcomes with Etanercept have been variable in pulmonary sarcoidosis (35), and other cytokines, such as IFN-γ may play more central roles in disease pathogenesis (36, 37). While our data do not support Etanercept monotherapy, they do not exclude potential efficacy of TNF-α inhibition in combination regimens or with alternative TNF-α-blocking agents (2, 3, 36).

The main limitation of this study is that it primarily focused on the acute inflammatory phase of AIM and the short-term effects of treatment. Longer-term follow-up is required to assess the impact of EC2319 on cardiac remodeling and sustained cardiac function, as fibrosis and ventricular dysfunction typically develop at later stages. Additionally, further studies are warranted to evaluate the long-term safety profile and biodistribution of EC2319, considering its mechanism of uptake via FR-mediated endocytosis.

In conclusion, FR-targeted antifolate therapy with EC2319 significantly reduces myocardial inflammation in a rat model of AIM, underscoring the pathogenic role of FR-β-expressing macrophages. [^18^F]FOL PET imaging provides a non-invasive method to detect myocardial inflammation and monitor treatment response, supporting its potential clinical utility for guiding and assessing immunosuppressive therapy in myocarditis.

## Methods

### Sex as a biological variable

Only male Lewis rats were used in this study. The use of a single sex was based on the well-established male predominance and reproducibility of the autoimmune myocarditis (AIM) model in Lewis rats (38). Male rats exhibit a consistent disease onset and severity following cardiac myosin immunization, which facilitates reduction of biological variability and allows clearer interpretation of treatment effects (38, 39). Female Lewis rats have been reported to show lower and more variable susceptibility to AIM, likely due to hormonal modulation of immune responses (38, 39). Therefore, inclusion of males only was scientifically justified for model consistency. The findings are expected to be relevant to both sexes, as the underlying immune mechanisms targeted by EC2319 are not sex-specific.

### Animal model and study protocol

AIM was induced in male Lewis rats by subcutaneous injections of 5 mg/mL pig cardiac myosin (M0531; Sigma Aldrich) in an equal volume of complete Freund’s adjuvant supplemented with Mycobacterium tuberculosis (1 mg/mL, F5881; Sigma Aldrich) into the hock of the left foot twice (on days 0 and 7) as previously described (10). To enhance immunization, the rats also received an intraperitoneal injection of 250 ng/mL pertussis toxin (P2980; Sigma Aldrich). Immunizations were done under isoflurane anesthesia, with post-immunization analgesia (buprenorphine 0.03 mg/kg, Richter Pharma AG, Wels, Austria; twice daily for 2 days).

Figure 5 shows the study protocol and number of rats involved. First, in the prevention sub-study, we evaluated the effect of EC2319 administered during immunization on the development of AIM. Starting on the day of the first immunization (on day 0), 24 rats were randomized to receive subcutaneous injections of EC2319 (750 nmol/kg body weight; Endocyte, West Lafayette, IN, USA) or saline in the nuchal area twice a week for three weeks (6 doses/animal). The animal weights were monitored weekly to track weight changes. On day 21, corresponding to the time of most active cardiac inflammation (10), rats were sacrificed for histological analyses of myocardial inflammation. LV function was assessed by echocardiography one day before the first immunization (on day -1) and at the end of intervention (day 20).

**Figure 5.**
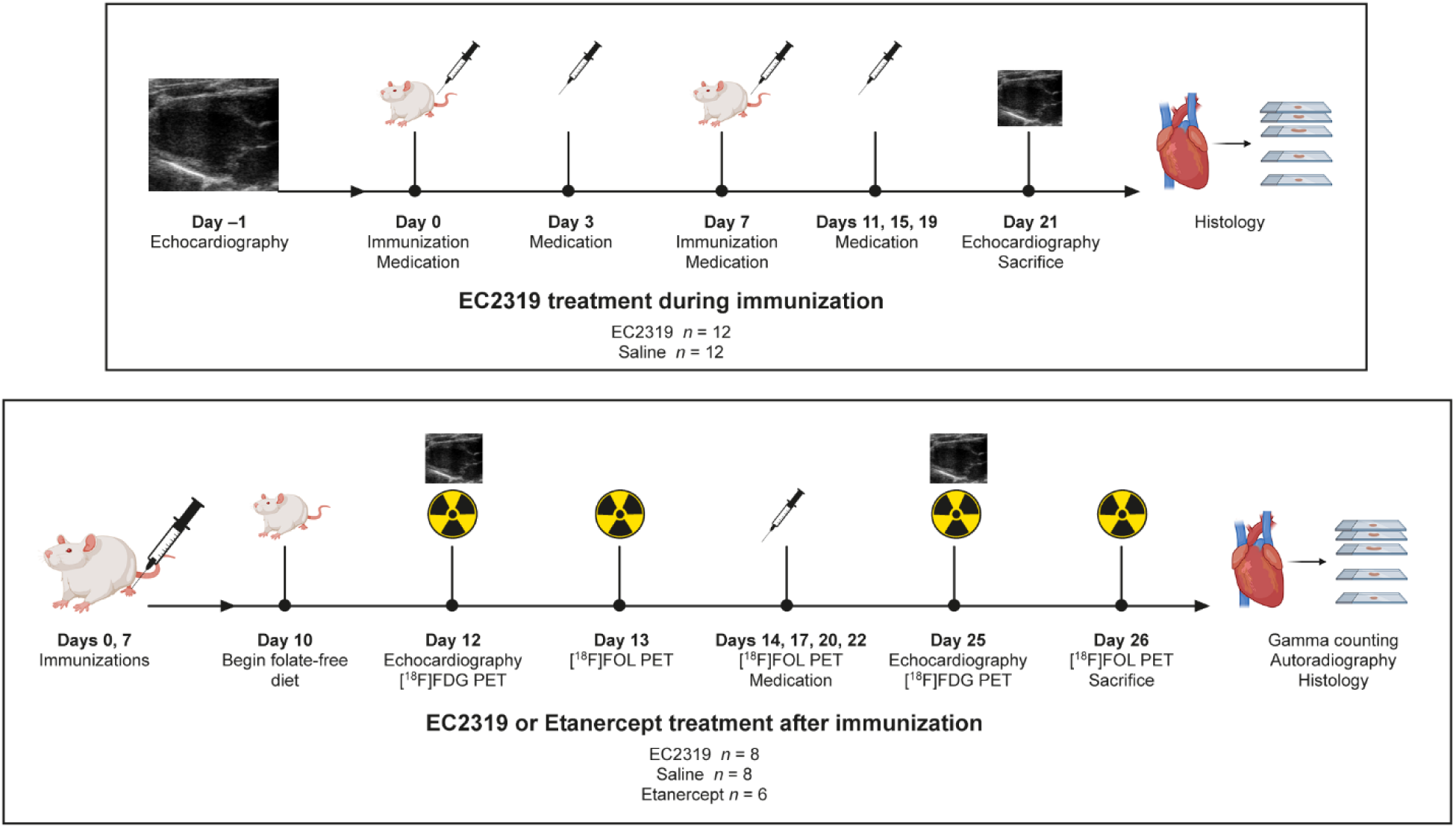
Study design (created using BioRender.com).

Next, in the intervention sub-study, we evaluated anti-inflammatory effects of EC2319 during active myocarditis in 46 rats. After completing immunizations on days 0 and 7, rats were placed on a folate-free diet (5T0F:57W5; Testdiet, St. Louis, MO, USA) on day 10 and underwent [^18^F]FOL PET on day 13 in order to detect active myocardial inflammation. Nine rats were lost, either shortly after immunization due to severe illness with respiratory distress refractory to supportive care, or during isoflurane anesthesia for the first echocardiography. Fifteen rats without active myocardial inflammation were excluded. The remaining 22 rats with focally increased myocardial [^18^F]FOL uptake indicating active myocarditis were then randomly divided into three groups. The first group (*n* = 8) received subcutaneous administration of EC2319 (750 nmol/kg body weight), the second group (*n* = 8) received subcutaneous saline, and the third group (*n* = 6) received subcutaneous Etanercept, a fully humanized recombinant TNF receptor (p75)-Fc fusion protein (10 mg/kg body weight; Immunex Corporation, Thousand Oaks, CA, USA) on days 14, 17, 20 and 22 (4 doses/animal). Four days after the last dose of EC2319, saline or Etanercept (on day 26), rats underwent second [^18^F]FOL PET scans and were sacrificed for ex vivo gamma counting of excised tissues, autoradiography of cardiac tissue sections, and histological analyses of myocardial inflammation. [^18^F]FDG PET was performed one day before each [^18^F]FOL PET scan (on days 12 and 25) as a visual reference of the myocardium. LV function was assessed by echocardiography before treatment (on day 12) and at the end of study (on day 25).

### Radiotracers

[^18^F]FDG was prepared using standard FASTLab cassettes (19). [^18^F]FOL was synthesized following established procedures (10, 16). The total radiosynthesis time ranged from 82 to 115 minutes, beginning at the end of bombardment. The resulting radiochemical purity exceeded 95%, with a molar activity of 1218.9 ± 184.5 GBq/μmol (*n* = 3). The decay-corrected radiochemical yields were 12.5 ± 6.6%. The in vivo stability of [^18^F]FOL was assessed previously (10), using blood samples obtained 80 minutes post-injection from three control rats and four immunized rats. The fraction of radioactivity corresponding to the intact tracer in plasma was 91 ± 2.3%.

### In vivo PET/CT imaging

In vivo imaging of [^18^F]FOL was conducted on rats anesthetized with isoflurane (induction 4.5-5%, maintenance 1.5-2.5%), using a small-animal PET/CT scanner (Inveon Multimodality; Siemens Medical Solutions) as outlined in previous studies (10). The rats received an injection of 46.7 ± 8.2 MBq of [^18^F]FOL via the tail vein, followed by a 20-minute static PET scan starting 20 minutes post-injection.

To visualize the myocardium, a separate 10-minute static PET scan using [^18^F]FDG (33.3 ± 6.4 MBq) was performed on the day before the [^18^F]FOL study, beginning 20 minutes after injection.

Using Carimas 2.10 software (Turku PET Centre, Turku, Finland), polar maps of [^18^F]FOL uptake in the LV myocardium were created as previously described (18). Myocardial contours were created in [^18^F]FDG images, which were then copied to co-registered [^18^F]FOL data. The polar maps were then generated with matching orientation and sampling points 20-40 minutes post-injection showing [^18^F]FOL as standardized uptake value (SUV) in 17 segments. To assess maximum intensity of inflammation, we measured uptake of [^18^F]FOL in segment with the highest SUV (SUV_max_). To assess global intensity and extent of myocardial inflammation, we measured mean [^18^F]FOL SUV (SUV_mean_) of the whole LV myocardium. To specifically assess the effect of treatments on myocardium with intense inflammation before therapy, we measured [^18^F]FOL SUV at the end of study in the segment with most intense inflammation before treatment (SUV_most inflamed segment_).

### Biodistribution analysis, autoradiography, histology, and immunostaining

Following PET/CT imaging, approximately 50 minutes post-injection of [^18^F]FOL, blood was collected via cardiac puncture, and the rats were euthanized by cervical dislocation. The hearts were rinsed with saline to remove excess blood and then excised. For biodistribution analysis, samples were taken from the heart, blood, plasma, pancreas, spleen, kidneys, intestine (without content), liver, lymph nodes, muscle, thymus, white adipose tissue, brown adipose tissue, and urine. The total radioactivity of these samples was measured using a gamma counter (Triathler 3’’, Hidex Oy, Turku, Finland). The radioactivity values were normalized for the injected dose per animal weight, decay, and tissue sample weight, and the results were expressed as SUV.

In the prevention sub-study, excised hearts were fixed in 10% formaldehyde, embedded in paraffin and cut into serial transverse 4 μm sections at 1 mm intervals from base to apex for histology and immunostainings.

In the intervention sub-study, the excised hearts were frozen in cooled isopentane and sectioned into serial 20-μm and 8-μm transverse cryosections at 1 mm intervals from apex to base for autoradiography and histological analyses. Autoradiography was performed according to previously described methods (10). For general histology, 20-μm cryosections were stained with hematoxylin and eosin (H&E), scanned with a digital slide scanner (Pannoramic 250 Flash, 3DHistech Ltd., Budapest, Hungary), and superimposed with autoradiographs. For immunohistochemistry, adjacent serial 8-μm cryosections were stained with monoclonal mouse anti-rat CD68 antibody (1:1000, MCA341GA, Bio-Rad) to identify macrophages. Additionally, MRC-1 (1:5000 dilution; Abcam, Cambridge, UK), and iNOS (1:500 dilution; Abcam) antibodies were used in three rats per treatment group, to detect anti-inflammatory and pro-inflammatory macrophages, respectively. To further characterize macrophage subsets in inflammatory lesions, double immunofluorescence staining was performed on cryosections. Sections were first stained with anti-CD68 antibody (1:500, clone ED1, MCA341GA, Bio-Rad, Oxford, UK), detected using goat anti-mouse IgG (recognizing both heavy and light chains) Alexa Fluor™ 488 (1:200, A11017, Invitrogen, Carlsbad, CA, USA), and subsequently stained with either anti-iNOS antibody (1:500, ab15323, Abcam, Cambridge, UK) or anti-MRC1 (CD206) antibody (1:5000, ab64693, Abcam, Cambridge, UK), both detected using goat anti-rabbit IgG (recognizing both heavy and light chains) Alexa Fluor™ 594 (1:500, A11037, Invitrogen, Carlsbad, CA, USA).

Furthermore, Masson trichrome staining was performed to study fibrosis. Immunostaining and fibrosis were quantified as stained area fraction (%) in the whole heart (CD68 and fibrosis) or within CD68-positive lesions (MRC-1 and iNOS) using ImageJ software (Fiji, National Institutes of Health, Bethesda, MD, USA).

### Echocardiography

Echocardiography was performed in isoflurane anesthetized rats (induction 4.5-5%, maintenance 1.5-2.5%) using a dedicated small animal Doppler ultrasound device (Vevo 2100, VisualSonics, Inc., Toronto, ON, Canada) with a linear 13-24 MHz (MS250) transducer as previously described (19). The LV end diastolic and end systolic volumes (LVEDV and LVESV, respectively) and LV ejection fraction (LVEF) were measured in parasternal long-axis 2D images.

### Statistical analysis

Continuous data are presented as mean ± SD unless specified otherwise. Normality distribution assumption was checked with the Shapiro–Wilk test, and the assumption of variance equality was assessed using Levene’s test. Statistical comparisons of normally distributed data were performed using paired Student’s t-tests to compare results before and after treatment, and unpaired Student’s t-tests or one-way ANOVA followed by post hoc tests: Tukey’s test (for homogeneous variances) and Games–Howell test (for non-homogeneous variances) to compare different treatments. Non-normally distributed continuous data are presented as median (interquartile range, IQR) and compared using the Kruskal–Wallis test, followed by pairwise comparisons with Bonferroni-adjusted p-values. Spearman correlation coefficient was used to analyze linear correlations between two continuous variables. All analyses were performed using SPSS version 29.0.2.0 and GraphPad Prism version 6, with a P value < 0.05 considered statistically significant.

## Study approval

All animal experiments were approved by the Finnish national Project Authorization Board (license ESAVI-37123-2020) and complied with the EU Directive 2010/63/EU.

## Data availability

This research was conducted in full compliance with all relevant ethical regulations. All data supporting the findings have been securely stored in the Turku PET Centre, Turku University Hospital database. Data may be made available upon reasonable request through a proposal-based process. Requests should be directed to the corresponding author, A.S.

## Author contributions

A.J. and E.A.H. contributed equally to the design and execution of experiments, data analysis and interpretation, and drafting the original manuscript. The order of co–first authors was determined by mutual agreement between the authors. W.N., J.V., H.L., J.P., A.P., X.Z., J.R., and X.G.L. participated in experiments, data analysis, and critical revisions of the manuscript. M.S., P.S.L., and Y.J.L. provided the drug candidate and radiotracer and contributed to critical revisions. J.K. oversaw project administration and provided critical revisions. A.R. and A.S. were responsible for conceptualization, study design, funding acquisition, and supervision.

## Acknowledgments

The authors thank Aake Honkaniemi and Marko Vehmanen for assistance in PET studies, Marja-Riitta Kajaala and Erica Nyman (Histology Core, Institute of Biomedicine, University of Turku) for tissue sectioning and immunohistochemical staining, and Timo Kattelus for assistance in figure preparation.

