## Supplemental File for "Targeting Macrophage Folate Receptor-β for Diagnosis and Treatment of Autoimmune Myocarditis"

### Supplemental Material

**Supplemental Table 1.** Ex vivo biodistribution of [$^{18}$F]FOL at 50 min post-injection.

| | EC2319 | Etanercept | Saline | *P*-value |
|---|---|---|---|---|
| **Blood** | 0.09 ± 0.01 | 0.03 ± 0.001 | 0.05 ± 0.005 | < 0.001* |
| **Brain** | 0.06 ± 0.01 | 0.04 ± 0.004 | 0.06 ± 0.007 | 0.300 |
| **Brown adipose tissue** | 0.70 ± 0.05 | 0.80 ± 0.07 | 0.90 ± 0.05 | 0.100 |
| **Heart** | 1.10 ± 0.10 | 3.60 ± 0.60 | 2.70 ± 0.40 | < 0.001* |
| **Small intestine without content** | 2.20 ± 0.10 | 4.10 ± 0.10 | 3.40 ± 0.60 | 0.070 |
| **Kidney** | 41.10 ± 2.60 | 17.50 ± 0.80 | 24.90 ± 2.70 | < 0.001* |
| **Liver** | 1.00 ± 0.08 | 4.00 ± 1.30 | 3.10 ± 0.70 | 0.030* |
| **Lung** | 0.80 ± 0.10 | 1.30 ± 0.10 | 1.20 ± 0.20 | 0.060 |
| **Lymph node** | 4.20 ± 0.30 | 4.10 ± 0.20 | 4.30 ± 0.10 | 0.600 |
| **Muscle** | 0.40 ± 0.03 | 0.40 ± 0.01 | 0.50 ± 0.09 | 0.300 |
| **Pancreas** | 1.10 ± 0.10 | 1.90 ± 0.30 | 5.20 ± 3.70 | 0.300 |
| **Pericardium** | 3.70 ± 0.30 | 0.03 ± 0.001 | 5.70 ± 1.00 | 0.200 |
| **Skull bone** | 0.20 ± 0.03 | 0.80 ± 0.08 | 0.40 ± 0.10 | 0.009* |
| **Spleen** | 2.60 ± 0.30 | 9.80 ± 0.60 | 5.90 ± 1.10 | < 0.001* |
| **Thymus** | 1.60 ± 0.30 | 3.20 ± 0.80 | 2.40 ± 0.20 | 0.050* |
| **Urine** | 11.90 ± 2.30 | 7.60 ± 1.00 | 12.40 ± 2.50 | 0.400 |
| **White adipose tissue** | 0.40 ± 0.05 | 0.40 ± 0.05 | 0.40 ± 0.07 | 0.300 |

The results are expressed as SUVs (mean ± SD). Statistical analysis done with one-way ANOVA.
